# Defining a Global Map of Functional Group Based 3D Ligand-binding Motifs

**DOI:** 10.1101/2020.09.27.315762

**Authors:** Liu Yang, Wei He, Yuehui Yun, Yongxiang Gao, Zhongliang Zhu, Maikun Teng, Zhi Liang, Liwen Niu

## Abstract

Uncovering conserved 3D protein-ligand binding patterns at the basis of functional groups (FGs) shared by a variety of small molecules can greatly expand our knowledge of protein-ligand interactions. Despite that conserved binding patterns for a few commonly used FGs have been reported in the literature, large-scale identification and evaluation of FG-based 3D binding motifs are still lacking. Here, we developed AFTME, an alignment-free method for automatic mapping of 3D motifs to different FGs of a specific ligand through two-dimensional clustering. Applying our method to 233 nature-existing ligands, we defined 481 FG-binding motifs that are highly conserved across different ligand-binding pockets. Systematic analysis further reveals four main classes of binding motifs corresponding to distinct sets of FGs. Combinations of FG-binding motifs facilitate proteins to bind a wide spectrum of ligands with various binding affinities. Finally, we showed that these general binding patterns are also applicable to target-drug interactions, providing new insights into structure-based drug design.

## Background

Protein-ligand interactions play fundamental roles in many important cellular functions, including small molecule metabolism, enzymatic catalysis, signal transduction and regulation. Comprehensive knowledge of protein-ligand interactions can not only provide important insights into biological functions of ligand-binding proteins but also greatly benefit drug discovery and development (Loewenstein et al., 2009; Paul et al., 2010). Many proteins that don’t display overall sequence or structure similarities may share similar local 3D structures and can bind to same or similar ligands, thus identifying conserved binding patterns across different ligand-binding proteins at 3D level could facilitate a better understanding of protein-ligand recognition (Abrusan and Marsh, 2018; Du et al., 2016; Persson et al., 2006).

With the rapid accumulation of experimentally determined structures of protein-ligand complexes, it became possible for large-scale identification of conserved 3D binding motifs using computational approaches (Kinjo and Nakamura, 2009; Ribeiro et al., 2020). Current methods based on structural comparison or alignment of protein pockets have identified many well-defined 3D motifs that are conserved across different protein pockets and widely used for protein function annotation, pockets classification and ligand-binding prediction (Gao and Skolnick, 2013; Hoffmann et al., 2010; Hwang et al., 2017; Pires et al., 2013; Pu et al., 2019; Yeturu and Chandra, 2008). However, these ligand-based 3D binding patterns are not applicable to large fraction of ligands especially small molecular drugs due to the lacking of reference 3D protein-ligand structures.

Despite the functional and structural diversity of different protein-binding ligands, many of them share same or similar functional groups (FGs) that mediate the interactions with the target proteins. Therefore, identification of conserved 3D binding motifs for FGs shared by different small molecules may extend our understanding of protein-ligand interactions to higher resolution and broader scope (Guvench, 2016). Previous studies have shown that conserved 3D motifs do exist in proteins binding different ligands with the same FG. For example, the phosphate-binding loop (P-loop) motif (Saraste et al., 1990; Via et al., 2000), in which the residues are highly conserved in terms of amino acid types as well as spatial positions among diverse phosphate-binding proteins. Conserved 3D motifs were also reported for other FGs such as adenine ring (Denessiouk et al., 2001; Narunsky et al., 2020; Nebel et al., 2007), heme group (Ferousi et al., 2019; Zubieta et al., 2007) and prosthetic groups (Nebel, 2006). However, these motifs were either uncovered through manual analysis of a small set of protein structures by an expert (e.g. a crystallographer) or through structural alignment of proteins that bind FGs with rigid structures, which are subjected to limited FG types and/or biased datasets of 3D structures. Computational methods for automatic extraction of 3D binding motifs for a variety of FGs in large scale are still lacking.

To systematically identify and evaluate 3D binding motifs at FG level, we developed AFTME, an alignment-free method that automatically maps functional atoms to different FGs from a set of protein pockets binding the same ligand using two-dimensional clustering approach. We applied our method to 233 natural ligands with abundant 3D protein-ligand structures and built an encyclopedia of 481 binding motifs for 160 different FGs, providing valuable resources for elucidating the mechanism of protein-ligand interactions as well as uncovering new rules for structure-based drug design.

## Results

### 1. AFTME enables automatic extraction of FG-based 3D binding motifs

We have developed AFTME, a computational method to dissect protein pockets binding a specific ligand into sectors that interact with different functional groups (FGs). The basic assumption of this method is simple: if conserved binding pattern for a specific FG exists, the pattern-forming atoms should be spatially proximal to the corresponding FG and frequently co-appear, thus can be detected through clustering analysis of functional atoms from diverse protein pockets binding the same ligand. Fig.1A outlines the major steps of the method. (1) Given a set of protein pockets binding the same ligand, AFTME first parses all the functional atoms (FAs) (He et al., 2015) that are considered to interact with the ligand atoms (LAs). (2) Then a distance matrix is constructed that evaluates the spatial distances between FAs and LAs. (3) Based on the distance matrix, a two-dimensional clustering algorithm is performed, through which LAs are clustered into different FGs at the first dimension and FAs are clustered into corresponding FG-binding motifs. (4) Each identified binding motif can be represented as a vector according to its chemical composition, which facilitate further analysis. The detailed description of each step is presented in Methods and Materials section.

Considering the abundance of studies on ATP-binding proteins, we first applied AFTME to a set of ATP-binding proteins as a proof of concept. As shown in Fig. 1B, our method identified three FA clusters or binding motifs corresponding to triphosphate group, ribose and adenine, respectively. The triphosphate-binding motif (M1) mainly consists of hydrophilic atoms from polar amino acids like ARG, LYS, SER, etc., and the adenine-binding motif (M2) is enriched by atoms from hydrophobic and aromatic amino acids including LEU, VAL, PHE, etc., whereas the ribose-binding motif (M3) contains both hydrophobic and hydrophilic amino acids (Fig. 1C). Further investigation of these binding motifs indicated that the AFTME identified FG-binding motifs are biological meaningful units. For instance, among the hydrophilic residues (rendered in red in Fig. 1D and Fig. S1) interacting with the triphosphate group, LYS and SER are both well-known conserved residues in the P-loop, a common motif for phosphate-binding in ATP-and GTP-binding proteins, which is typically composed of a glycine-rich sequence followed by a conserved lysine and a serine or threonine (Saraste et al., 1990). It was found that the hydrophobic and/or aromatic residues (rendered in green in Fig. 1D and Fig. S1) making up the adenine-binding motif interact with adenine ring through C-H–π and/or π–π interactions. Notably, Moodie *et al*. described the recognition of adenine by proteins in terms of a fuzzy recognition template based on a sandwich-like structure formed by hydrophobic residues (Moodie et al., 1996). Denessiouk *et al. also* found that bulky hydrophobic residues can form a hydrophobic area by interacting with the adenine base (Denessiouk and Johnson, 2000). The A-loop motif, which includes aromatic residues forming π–π interactions with adenine ring was also reported (Ambudkar et al., 2006). These findings are largely consistent with the AFTME identified adenine-binding motif. Although no well-defined motifs corresponding to the ribose-binding motif have been reported yet, it makes sense that hydrophobic/aromatic residues of the ribose-binding motif interact with five-carbon ring while hydrophilic residues interact with the extended hydroxyl groups through polar interactions.

**Figure 1.**
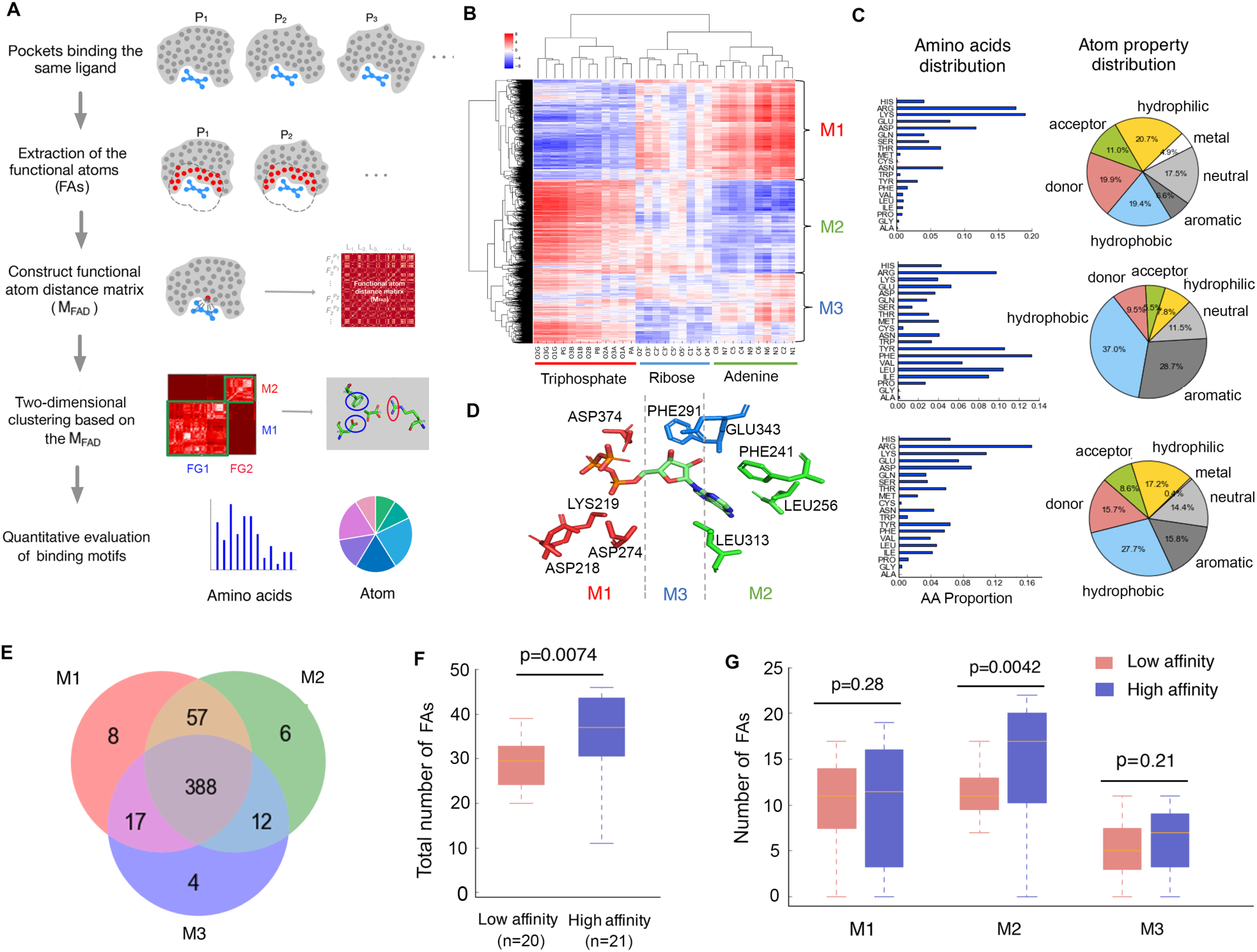
Workflow of AFTME and its application to ATP-binding proteins. (A) A schematic view of major steps of the AFTME method. (B) Two-dimensional clustering of the distance matrix for ATP-binding pockets. The vertical and horizontal axes correspond to FAs and LAs, respectively. The color encodes the distance between an FA and an LA. Three LA clusters corresponding to the triphosphate group, ribose and adenine ring of ATP were identified, respectively. Three FA clusters or binding motifs (M1, M2 and M3) corresponding to the above three FGs were also obtained simultaneously. (C) Distribution of amino acids and atom properties for M1(top), M2(middle) and M3 (bottom). (D) An example (PDB: 1VJC) showing the spatial distribution of amino acids within each identified FG-binding motif. (E) Different ATP-binding proteins use different combinations of FG-binding motifs M1, M2 and M3. (F) ATP-binding pockets with high affinity have significantly more FAs than those with low affinity. (G) Comparison of the FA numbers in FG-binding motifs of ATP-binding pockets with high and low affinity. The center line, bounds of box and whiskers represent the median, interquartile range and 1.5 times interquartile range, respectively. The p-values were calculated using Manney-Whitney test.

We then set out to explore different roles played by the three identified motifs in the ATP-binding process. As we can see from the Venn diagram in Fig. 1E, among the 492 ATP-binding pockets in the dataset, a majority (388, 75.9%) contain all the three binding motifs. Nevertheless, 86 (16.8%) pockets get two of them, among which 57 (66.3%) carry the phosphate-binding (M1) and adenine-binding (M2) motifs, indicating that combination of two motifs, especially M1 and M2, is sufficient for ATP binding. Besides, we also noticed some cases in which only one binding motif together with one or more metal ions exists (Fig. S2), indicating that metal ions may greatly affect the global binding profile. Next, we asked how different FG-binding motifs contribute to the binding affinity. All the ATP-binding proteins with experimental affinity data available were collected and sorted from high to low affinities (Table S1). In general, protein pockets in high-affinity (top 1/3) group contain more FAs than those in low-affinity (bottom 1/3) group (Fig. 1F, p=7.4e-03, Mann-Whitney test). Interestingly, when looking into an individual binding motif, only adenine-binding motif (M2) shows significant increment of FA numbers in high-affinity pockets (Fig. 1G, p=4.2e-03, Mann-Whitney test), suggesting that increase of hydrophobic interactions with the adenine ring makes major contributions to higher ATP binding affinity.

Taken together, the above results demonstrate the ability of AFTME to decompose ligand-binding sites into biological meaningful motifs, which are spatially well-defined to interact with different FGs and contribute unequally to protein-ligand binding affinity.

### 2. FG-based binding motifs are reused among different ligand-binding proteins

To see whether FG-based 3D binding motifs identified by our method are reused across different ligand-binding proteins, we applied AFTME to a few ligands sharing the same FGs with ATP including ADP, AMP, GTP and UTP. Fine-mapped binding motifs for adenine and ribose were obtained from ADP-and AMP-binding proteins, and that for triphosphate was from GTP-and UTP-binding proteins (Fig. S3A-3D). We found that the chemical compositions of motifs binding the same FG are highly consistent although they were extracted from proteins binding different ligands (Fig. 2A-2C). For adenine and ribose, the binding motifs are universal among ATP-, ADP-and AMP-binding pockets and have very similar distribution of amino acid types and atom categories (Fig. 2A, 2B). Similarly, triphosphate-binding motifs extracted from GTP-and UTP-binding proteins also show consistent makeup with ATP-derived motifs (Fig. 2C).

**Fig 2.**
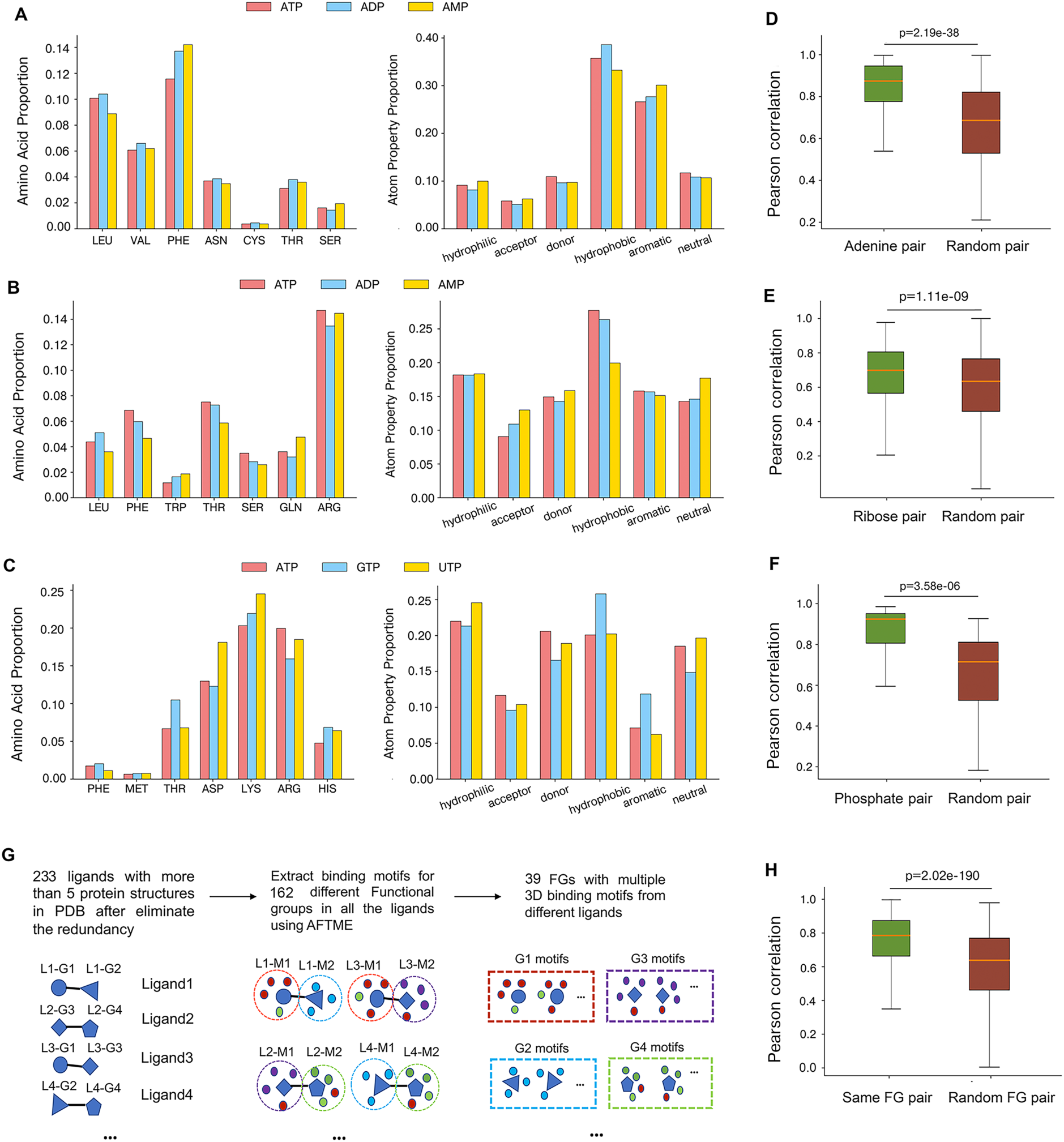
Conservation of FG-based binding motifs. (A-C) The chemical compositions of (A) adenine-, (B) ribose-and (C) triphosphate-binding motifs identified from proteins binding different ligands are similar. Adenine-and ribose-binding motifs are extracted from ATP-, ADP- and AMP-binding proteins, and triphosphate-binding motifs are obtained from proteins binding ATP, GTP and UTP, respectively. (D-F) Pairs of (D) adenine-, (E) ribose- and (F) triphosphate-binding motifs show significantly higher composition correlation than motif pairs binding random FGs. (G) A schematic view of large-scale identification of FG-based binding motifs using AFTME, the L, G and M represent the Ligand, the FG and the Motif, respectively. (H) Correlation analysis of the 481 identified FG-binding motifs indicates that motifs binding the same FG are highly consistent in their composition. The center line, bounds of box and whiskers represent the median, interquartile range and 1.5 times interquartile range, respectively. The p-values were calculated using paired t-test.

We then extended our evaluation to a wider range of ligands, among which 25 are adenine-containing, 38 are ribose-containing and 9 are triphosphate-containing. We described each FG-binding motif using a 26-dimensional vector representing the proportion of 20 types of amino acids and 6 categories of atoms, respectively (see Methods and Materials). The vector representation of the FG-binding motifs enables correlation analysis of chemical composition for any pair of motifs. It was found that adenine-binding motif pairs showed significantly higher correlations than random FG-binding motif pairs (Fig. 2D). Similar observations were obtained for ribose-binding (Fig. 2E) and triphosphate-binding (Fig. 2F) motif pairs, respectively. These results suggest high conservation of FG-binding motifs across a diversity of ligand-binding proteins.

Next, we performed a large-scale analysis of 3D FG-binding motifs for all the ligands with abundant 3D structures available. As shown in Fig. 2G, we first derived all the 3D protein-ligand structures from BioLiP database (Yang et al., 2013). Redundant proteins binding the same ligand that show over 50% sequence similarity were eliminated using cd-hit (Li and Godzik, 2006). 233 ligands with more than 5 structures available were kept for the following analysis. 481 FG-binding motifs corresponding to 160 unique FGs were identified using AFTME (Table S2), among which 39 FGs appeared in multiple (at least three) ligands. For each FG present in multiple ligands, a conservation score (CS) of the corresponding FG-binding motif was calculated as the average of pairwise Pearson’s correlation coefficients among all the identified motifs binding this specific FG. And a corresponding p value was also calculated using permutation test (see Methods and Materials). As shown in Table 1, most FG-binding motifs corresponding to FGs appeared in multiple ligands are highly conserved across different ligand-binding proteins (CS > 0.6, p < 0.05). Overall, two motifs binding the same FG show significantly higher composition correlations compared with two randomly selected motifs (Fig. 2H), confirming the high conservation of FG-binding motifs. These lines of evidences showed that AFTME can be applied to detect binding motifs for a diversity of FGs. Importantly, the identified binding motifs are highly conserved among different ligand-binding pockets, laying the foundations for expanding limited 3D motifs to a broader range of ligands that are not suitable for AFTME analysis (e.g. due to lack of structure data) but sharing same or similar FGs with applicable ligands.

**Table 1.** Conservation evaluation of binding motifs for multi-ligand FGs. The frequency reported the number of ligands containing the FG in the dataset, the conservation score was calculated as the average of pairwise Pearson’s correlations among all the FG-binding motifs for the specific FG, the p-value was calculated using permutation test.

| FG name | Frequency | Conservation score | p-value |
| --- | --- | --- | --- |
| Ribose | 38 | 0.665 | 1.00E-06 |
| Phosphate | 37 | 0.714 | 1.00E-06 |
| Carboxyl | 31 | 0.784 | 1.00E-06 |
| Glucose | 28 | 0.842 | 1.00E-06 |
| Adenine | 25 | 0.848 | 1.00E-06 |
| Pyrophosphate | 21 | 0.669 | 3.70E-05 |
| Guanine | 11 | 0.723 | 8.00E-06 |
| Uracil | 10 | 0.853 | 1.00E-06 |
| Acetic acid | 10 | 0.752 | 1.00E-06 |
| Triphosphate | 9 | 0.857 | 1.00E-06 |
| Propylamine | 8 | 0.713 | 4.46E-03 |
| Alanine | 8 | 0.856 | 1.00E-06 |
| 2-Deoxyribose | 8 | 0.718 | 2.17E-03 |
| Hydroxyethyl | 7 | 0.678 | 7.78E-02 |
| Acetamido group | 6 | 0.776 | 2.90E-04 |
| Glutamic acid | 6 | 0.7 | 4.36E-02 |
| Hexane group | 6 | 0.907 | 1.00E-06 |
| Hydroxymethyl | 6 | 0.74 | 4.55E-03 |
| Cytosine | 5 | 0.646 | 3.16E-01 |
| Pteridine rings | 5 | 0.773 | 3.26E-03 |
| Phenol | 5 | 0.88 | 1.00E-06 |
| Butyric acid | 5 | 0.723 | 4.05E-02 |
| Adenosine | 5 | 0.845 | 3.00E-06 |
| Glycine | 4 | 0.748 | 4.27E-02 |
| Ethylamine | 4 | 0.756 | 3.70E-02 |
| Para-amino benzoic acid | 4 | 0.692 | 1.73E-01 |
| Galactose | 4 | 0.92 | 1.00E-06 |
| Thymine | 4 | 0.959 | 1.00E-06 |
| Phenyl | 4 | 0.726 | 1.27E-01 |
| Glycolic acid | 4 | 0.741 | 5.21E-02 |
| Fructose | 4 | 0.759 | 4.16E-02 |
| Maltose | 4 | 0.819 | 2.14E-03 |
| Sulfo | 4 | 0.636 | 4.11E-01 |
| Glycerol | 3 | 0.869 | 3.77E-03 |
| Nicotinamide | 3 | 0.817 | 2.63E-02 |
| Propyl | 3 | 0.748 | 1.38E-01 |
| Ribose-3-phosphate | 3 | 0.933 | 4.30E-05 |
| Pyruvic acid | 3 | 0.821 | 2.35E-02 |
| Dimethylallyl | 3 | 0.799 | 4.41E-02 |

### 3. Towards an encyclopedia of 3D binding motifs for a diversity of FGs

Given that 481 binding motifs for 160 different FGs have been identified using our method, we asked whether there are general interaction patterns between the identified motifs and the FGs they bind. We found that all the binding motifs could be clustered into 4 classes based on their physicochemical properties using k-means (Figure S4, see Method and Materials), which are well separated in the t-SNE plot shown in Fig. 3A. Notably, the FG-binding motifs in different classes are featured with distinct physicochemical properties. The first class (red dots), denoted as the aromatic motif class, is enriched with atoms from aromatic amino acids like TRP, TYR and PHE. The second class (green dots), named the hydrophilic motif class, is mainly composed of hydrophilic, donor and acceptor atoms from polar amino acids such as ARG, LYS, ASP and GLU. The third class (blue dots), the mixed motif class, consists of both aromatic and hydrophilic atoms. The fourth class, named the hydrophobic motif class, is dominated by atoms from hydrophobic amino acids including LEU, ILE, VAL etc. (Fig. 3A).

**Fig 3.**
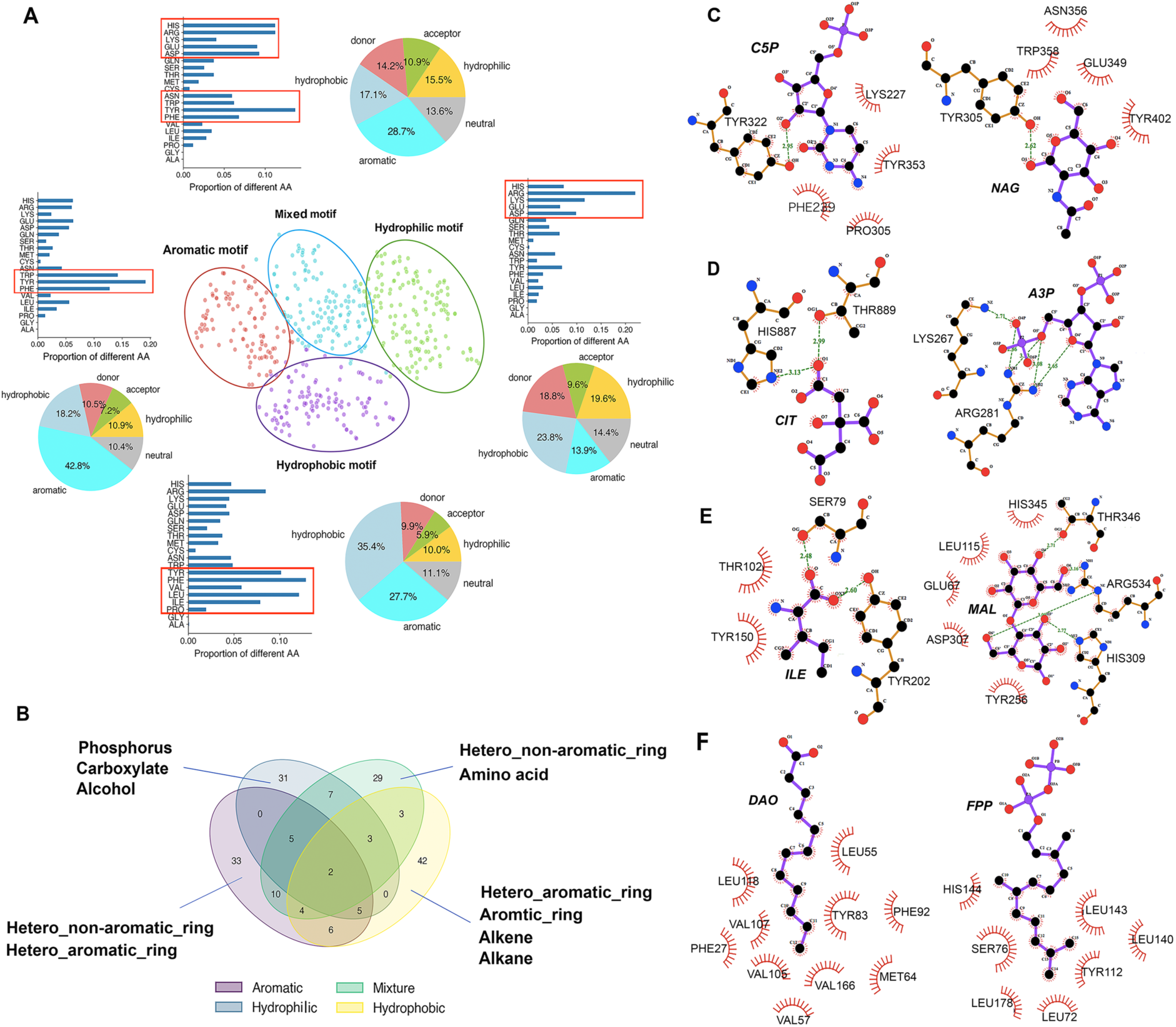
Clustering of FG-based binding motifs. (A) FG-binding motifs can be clustered into four well-separated classes, each of which has distinct distribution of amino acids (bar plot with the major amino acid types marked in red rectangular box) and atom types (pie plot). (B) Different motif classes show different FG-binding preferences, but some FGs are shared by motif classes. Dominant FG types for each motif class are denoted beside the Venn plot. (C-F) Examples of 2D interaction map between FGs and identified motifs. (C) The aromatic motifs identified for cytosine ring of cytidine-5’-monphosphate (PDB ID:4G5T, left) and glucose ring in N-acetyl-D-glucosamine (PDB ID: 6EN3, right). (D) The hydrophilic motifs identified for the carboxyl group of citric acid (PDB: 6FXI, left) and the phosphate group of adenosine-3’-5’-diphosphate (PDB ID: 1KAI, right). (E)The mixed motifs identified for amino acid isoleucine (PDB ID: 1Z17, left) and two glucose rings in maltose (PDB ID: 1AHP, right). (F)The hydrophobic motifs identified for the hexane group of lauric acid (PDB ID: 2OVD, left) and the farnesyl group in farnesyl diphosphate (PDB ID: 2E90, right). The 2D ligand-protein interactions were generated by LigPlot (Laskowski and Swindells, 2011).

Next, we looked into the correspondence between different motif classes and the FGs they bind. As shown in Figure 3B, most of the FGs are uniquely mapped to a single motif class, indicating that different classes of motifs have their specific binding preference for FGs. Although a variety of FGs are involved, we found some dominating FGs in each motif class (Table S3).

Among FGs that interact with the aromatic motifs, two types of FGs are in the majority, one is with aromatic ring and the other is with non-aromatic ring. The former type, exemplified with the cytosine ring of cytidine-5’-monphosphate (C5P), interacts with the aromatic ring of PHE and TYR through π-stacking (Fig. 3C left panel). The latter type, for example, the glucose ring in N-acetyl-D-glucosamine (NAG), of which the carbon atoms form hydrophobic interactions with the aromatic atoms of TYR/TRP (Fig.3C right panel).

In contrast, the hydrophilic motif class prefers to bind polar FGs through hydrogen bonds, among which carboxyl and phosphorus are the most prevalent ones. For instance, the carboxyl group in citric acid (CIT), forms N–H⋯O and O-H⋯O hydrogen bonds with N atom from imidazole of a HIS and O atom from hydroxyl of a THR, respectively (Fig.3D left panel). Four N–H⋯O hydrogen bonds are formed between O atoms of the phosphate group in adenosine-3’-5’-diphosphate (A3P) and N atoms of two basic amino acids (LYS and ARG) in the binding motif (Fig.3D right panel).

There are over 10 FGs engaged in both the mixed and the aromatic motif classes, most of which are non-aromatic sugar rings. In addition to hydrophobic interactions between the sugar ring and the aromatic ring which are frequently used in the aromatic motif class, the mixed motif class also contains hydrophilic amino acids that form hydrogen bonds with the extended-out hydroxyl groups (Fig. 3E, right panel). Besides, the mixed motif class is of high propensity to recognize an amino acid, of which the amide group interacts with the aromatic ring through amide-π stacking and the carboxyl group interacts with hydrophilic amino acids via hydrogen bonds, respectively (Fig. 3E left panel).

The hydrophobic motif class also shares a major type of FG, the aromatic hetero-ring, with the aromatic motif class. Instead of π–π interactions, the C-H–π interactions are the main driving force for hydrophobic-aromatic contacts. Another two major types of FGs involved in the hydrophobic motif class are alkene and alkane chains. As two examples showed in Figure 3F, the alkane (left) and the alkene (right) chains are well accommodated in protein pockets composed of hydrophobic residues.

Altogether, our systematic analysis suggested the existence of four classes of FG-binding motifs and their favored FGs. Deep investigations further revealed general interaction patterns between these functional motifs and the FGs they bind, thus build up a global map of 3D motif-FG interactions.

### 4. Motif combinations facilitate different modes of ligand-binding

Having identified the corresponding relations between motif classes and FGs, we then asked how motifs are combined to facilitate the binding of ligands that consist of different FGs. We found that the above four FG-binding motif classes are almost evenly distributed in protein pockets investigated in our analysis (Fig. 4A), suggesting that all the identified motif classes are commonly used and important for protein-ligand recognition.

**Fig 4.**
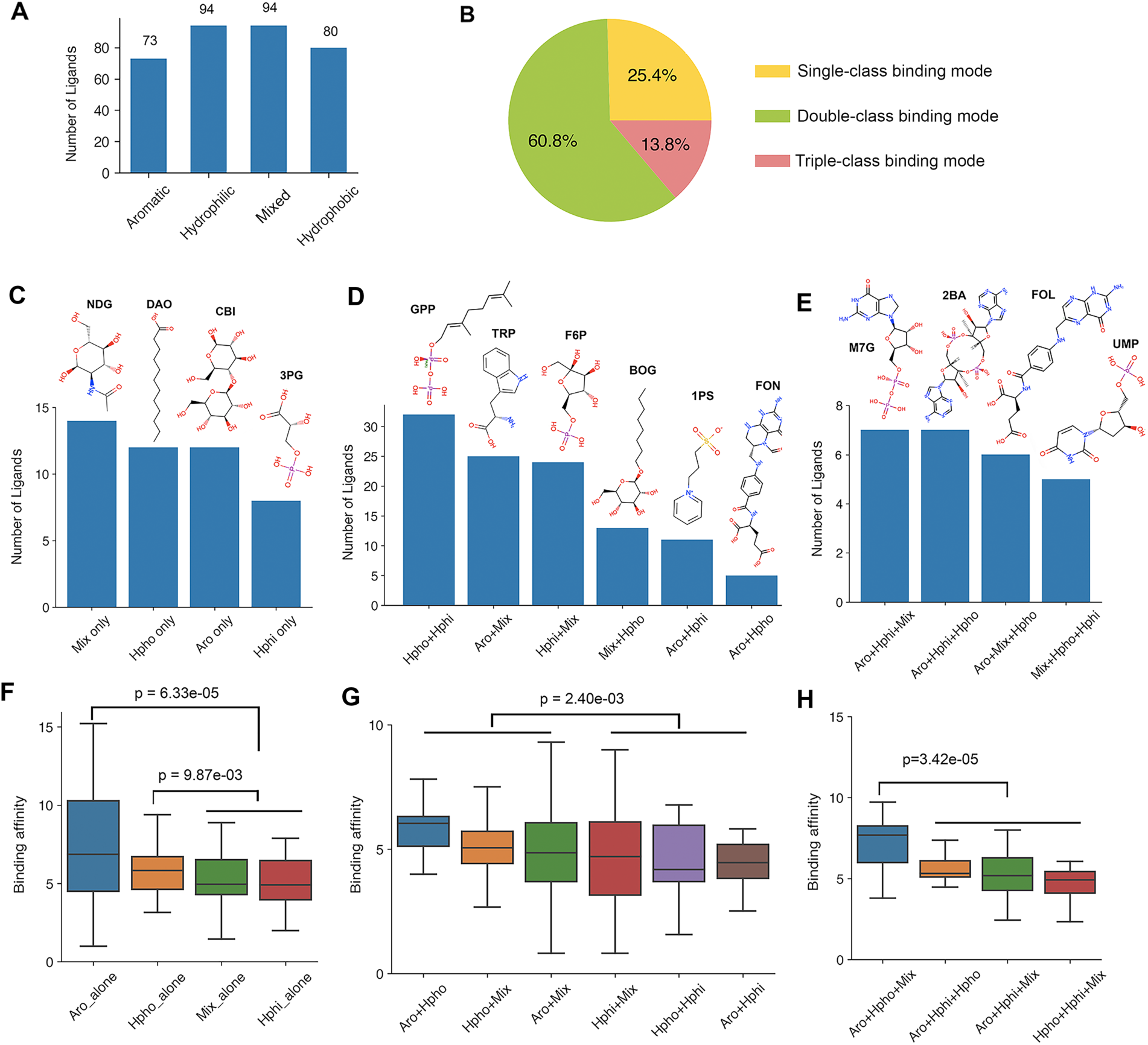
Combinations of FG-binding motif classes in a ligand-binding pocket. (A) Distribution of the four classes of FG-binding motifs in ligands. (B) Proportion of the three different combination modes for motif classes. (C-E) Distribution of different FG-binding motif combinations and examples of ligands involved in (C) single-class (D) double-class and (E) triple-class mode. (F-H) Ligand-binding affinity is affected by combination of FG-binding motif classes for the (F) single-class, (G) double-class and (H) triple-class combination modes. Mix, Hpho, Aro and Hphi refer to mixed-class, hydrophobic-class, aromatic-class and hydrophilic-class motifs, respectively. The center line, bounds of box and whiskers represent the median, interquartile range and 1.5 times interquartile range, respectively. The p-values were calculated using Manney-Whitney test.

After careful inspection of the identified binding motifs and their host ligand-binding pockets, we found three distinct combination modes for motif classes in protein pockets (Fig. 4B). (i) The single-class mode, which applies to nearly a quarter of investigated ligand-binding cases, combines only FG-binding motifs of the same class. (ii) The double-class mode, which goes for more than 60% of the cases, integrates two different classes of FG-binding motifs. (iii) The triple-class mode, which recognizes a smaller fraction of ligands, assembles three different classes of FG-binding motifs.

For the single-class mode, combinations of two mixed-class motifs are mostly observed, followed by hydrophobic-, aromatic- and hydrophilic-class motif combinations (Fig. 4C). For the double-class mode, there are 6 possible class-class combinations, among which the hydrophobic-hydrophilic combination applies to the greatest number of ligands, indicating a commonly used protein-ligand binding pattern in which the hydrophobic FG of the ligand interacts with a hydrophobic motif while another polar FG is oriented to a hydrophilic motif (Fig. 4D, Fig. S5). The triple-class mode also includes four different class-class combinations that are almost equally present for ligands they bind (Fig. 4E).

To gain further insights, we investigated in greater detail of the ligands involved in different combination modes (Table S4). Notably, combinations of different classes of FG-binding motifs facilitate the binding of a vast diversity of ligands composed of FGs that are well mapped to the corresponding FG-binding motif classes.

Among ligands involved in the single-class mode, we outlined three for examples (Fig. 4C, Fig. S6A). 2-acetamido-2-deoxy-alpha-D-glucopyranose (NDG) is composed of two mixed-motif favored FGs including an acetamide group and a glucose ring, both being bound by FG-binding motifs of the mixed class. Cellobiose (CBI) is a disaccharide consisting of two glucoses and binds to proteins with two aromatic motifs. 3-phosphoglyceric acid (3PG) has two polar FGs, a phosphate group and a glyceric acid group, which are recognized by two hydrophilic motifs.

A greater number and higher variety of ligands were witnessed in the double-class mode (Fig. 4D, Fig. S6B). For instances, geranyl diphosphate (GPP) which is complexed with proteins comprising a hydrophobic and a hydrophilic motif, contains a hydrophobic-preferred alkene and a hydrophilic-preferred diphosphate group. For fructose-6-phospahte (F6P) proteins achieve ligand-binding with an aromatic motif to the sugar ring and a hydrophilic motif to the phosphate group, respectively. Other ligands such as tryptophan (TRP, with an aromatic motif to indole and a mixed motif to alanine), B-octyl glucoside (BOG, with a hydrophobic motif to octyl and a mixed motif to glucose), 3-pyridinium-1-ylpropane-1-sulfonate (1PS, with an aromatic motif to pyridinium and a hydrophilic motif to sulfonate) all follow the general FG-motif interaction patterns we identified.

In the triple-class mode, the ligands have at least three FGs and thus are in relatively larger size (Fig. 4E, Fig. S6C). For examples, to bind 7N-methyl-8-hydroguanosine-5’-diphosphate (M7G), proteins adopt a binding pattern with an aromatic motif to dehydroalanine, a mixed motif to ribose and a hydrophilic motif to diphosphate. Similarly, folic acid (FOL) contains a pteridine ring, a benzoic group and a glutamic acid that are recognized by a hydrophobic, an aromatic and a mixed motif, respectively.

In the example of ATP, we already showed the unequal contribution of different FG-binding motifs to the binding affinity. Here, we sought to explore how combinations of different classes of FG-binding motifs will affect the ligand-binding affinity. Experimental binding affinity for all the protein-ligand pairs in our analysis were retrieved from the PDBbind database (Table S5) (Liu et al., 2015). In the single-class mode, both the aromatic and hydrophobic combinations show significantly higher affinity than the mixed and hydrophilic combinations, suggesting that general hydrophobic interactions including π–stacking, C-H–π interactions and interactions between two aliphatic carbons contribute more to high binding affinity than polar interactions such as hydrogen bonds and salt bridge (Fig. 4E). Consistently, in the other two combination modes, the more hydrophobic interactions are involved, the higher affinity the ligand-binding achieve. For example, combination of the hydrophobic and aromatic motifs is the most efficient binding pattern in the double-class mode. Moreover, non-hydrophilic combinations get significantly higher affinity compared to combinations with hydrophilic motifs for both the double-class (Fig. 4F, p=2.4e-03, Mann-Whitney test) and triple-class modes (Fig. 4G, p=3.42e-05, Mann-Whitney test). The results further supplemented our observation from the ATP-binding motifs (Fig. 1G) and confirmed the findings in the previous studies that hydrophobic interactions are a driving factor for the increased ligand efficiency (Ferreira de Freitas and Schapira, 2017; Young et al., 2007).

Together, these evidences showed that FG-binding motifs are building blocks of ligand-binding sites, and combinations of different classes of FG-binding motifs facilitate the proteins to bind a wide spectrum of ligands with various binding affinities.

### 5. Motif-FG binding patterns are applicable to target-drug interactions

We next asked whether the above motif-FG binding patterns derived from nature-existing protein-ligand complexes can also be applied to design target-drug interactions. To this end, we investigated in details of interactions between three well-defined drug targets and their small molecular inhibitors (Fig. S7A). The first case is the kinase domain of BRAF, which is the target of vemurafenib, an FDA-approved small molecular drug for the treatment of patients with metastatic melanoma with BRAF V600E mutation (Karoulia et al., 2016). Four FAs clusters (motifs) were identified based on their spatial distances to LAs, that are well separated and mapped to different FGs of vemurafenib in 3D structure (Fig. 5A, 5B, Fig. S7B). The first motif mainly consists of hydrophobic atoms and interacts with the chlorophenyl group through hydrophobic interactions. The second motif contains atoms from two aromatic amino acids (TRP and PHE) and contacts with the pyridinyl group through π–stackings. The third motif includes both hydrophobic and hydrophilic atoms engaged in the hydrophobic and polar interactions with the difluorophenyl-sulfonamide group. The fourth motif is dominated by hydrophobic atoms that interact with the propane group. The second case is the methyltransferase domain of DOT1L, which is the target of EPZ-5676, a small molecular drug in clinical trial for the treatment of adult acute leukemia (Stein et al., 2018). Using the same approach, we found a hydrophobic motif (M1), a mixed motif (M2) and a hydrophilic motif (M3) in the ligand-binding sites of DOT1L, which correspond to three different FGs of EPZ-5676, i.e. methyl-adenine, ribose and methionine, respectively (Fig. 5C, 5D, Fig. S7C). Lastly, we looked into the interactions between the main protease (Mpro) of COVID-19 and one of its potent inhibitor 11b (Dai et al., 2020). We observed three well-separated motifs in the ligand-binding sites of the target protein, which are located proximal to the pyrrolidine, fluorophenyl and indole-carboxamide groups (Fig. 5E, Fig. S7D). Notably, the interactions between the identified FG-binding motifs and the corresponding FGs also follow the general chemical rules: two hydrophobic motifs (M1 and M2) interact with two ring structures, the mixed motif (M3) interacts with the carboxamide (Fig. 5F).

**Fig. 5.**
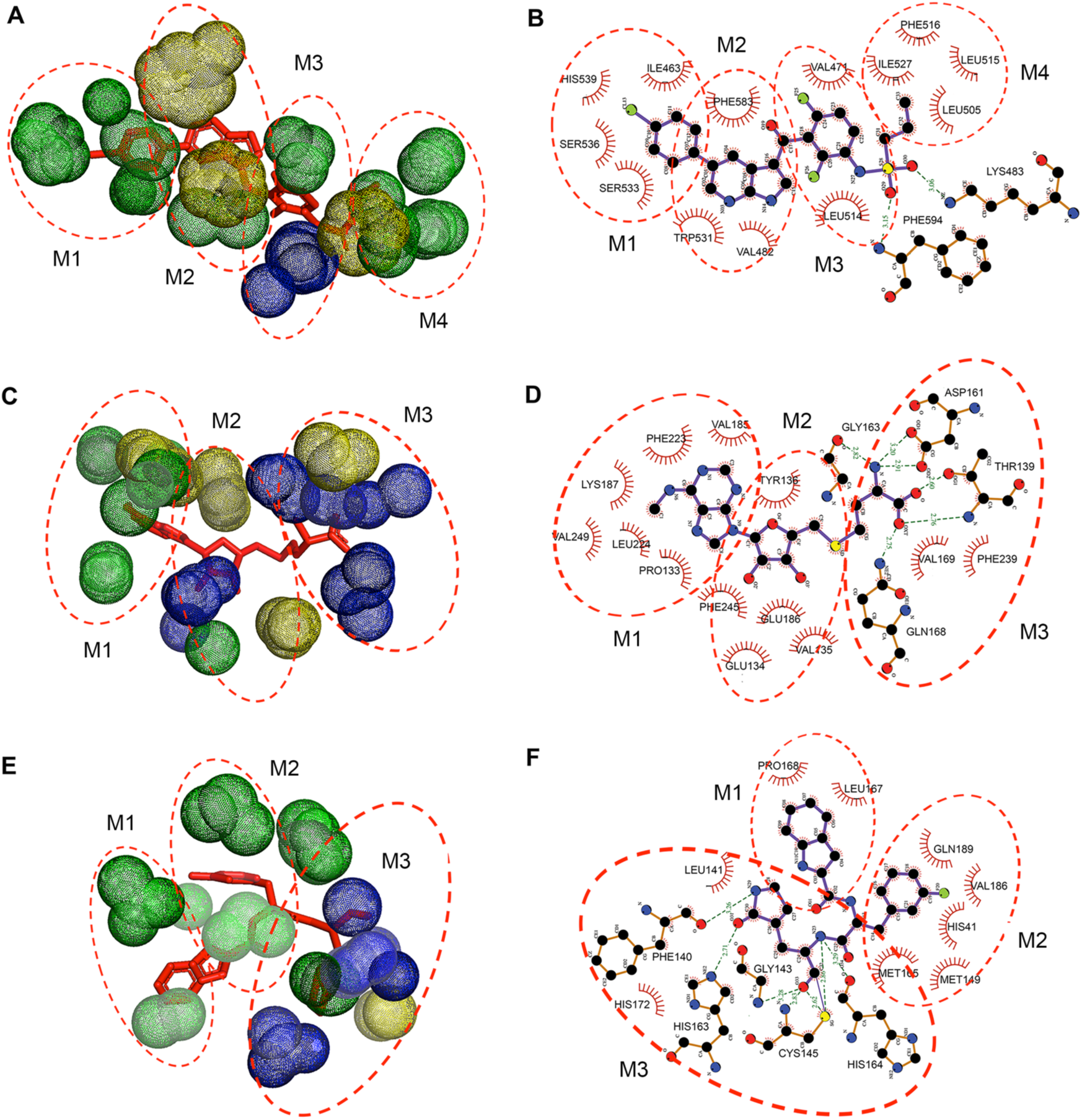
Three examples of motif-FG binding patterns in target-drug interactions. (A-F) 3D map showing the distribution of FG-binding motifs relative to different FGs of small molecular drugs (left) and the corresponding 2D ligand-protein interaction map (right) for (A-B) BRAF-vemurafenib complex (PDB ID: 4RZV), (C-D) DOT1L-EPZ-5676 complex (PDB ID: 3SR4), and (E-F) Mpro of COVID-19 in complex with 11b (PDB ID: 6M0K). Atoms with different physicochemical property are rendered in different colors: hydrophobic (green), polar (purple) and aromatic (yellow). The 2D ligand-protein interaction maps are generated by LigPlot (Laskowski and Swindells, 2011).

Together, these examples indicate that the FG-based functional motifs also appear in different drug targets and interact with the specific FGs following general motif-FG binding patterns. Thus, the global map of 3D FG-binding motifs could provide important insights and guidance for rational design of small molecular drugs.

## Discussion

A classical assumption in structural biology is that the 3D structure of a protein determines its molecular function. However, many proteins that don’t display overall sequence or structure similarities may share similar local 3D binding sites and can bind to same or similar ligands (Kahraman et al., 2007). Thus, identifying conserved 3D patterns/motifs across different ligand-binding proteins serve as an efficient way to learn and predict protein-ligand interactions. Computational methods that rely on multiple structure alignments or pairwise pocket comparisons have identified many conserved 3D binding patterns across different protein pockets binding same or similar ligands (Dukka, 2013). Despite the validity and usefulness of these ligand-based binding patterns, they mainly go for nature-existing ligands with abundant protein-ligand 3D structures, thus limiting their application scope. Here, we proposed AFTME, an alignment-free method for automatic identification of 3D binding motifs at the basis of FGs shared by different small molecules, which permits studying protein-ligand interactions in a wider scope and higher resolution. The application to ATP showed the feasibility and validity of our method to detect FG-based 3D binding motifs and confirmed the reusability of the motifs in different ligand-binding proteins. We further applied our method to 233 natural ligands and obtained 481 binding motifs for 160 unique FGs, providing useful resources for deep exploration of protein-ligand recognition.

Systematic investigation of FG-binding motifs identified by our method provides several important insights into protein-ligand interactions. First, ligand-binding sites of a protein can be dissected into independent sectors corresponding to different FGs of the binding ligand. These FG-based binding motifs are highly conserved among different ligand-binding pockets at both amino acid and atom level. Second, we found four classes of FG-binding motifs with distinct physicochemical properties and their own preference for FG binding. Moreover, the interactions between 3D motifs and FGs follow some general rules. For example, a hydrophobic motif is more likely to interact with a hydrophobic FG and a hydrophilic motif usually recognizes a polar FG. Third, following the general motif-FG recognition map, protein pockets consisting of different FG-binding motifs can bind a wide spectrum of ligands through different motif combination modes. Of note is that protein pockets with more hydrophobic motifs tend to gain higher binding affinity. Although the FG-binding motifs are mainly derived from protein structures binding natural ligands, we showed that these motifs can also be applied to different drug-target interactions. Therefore, we expected that the global map of motif-FG interactions, together with the current molecular docking (Trott and Olson, 2010) and/or molecular simulation (MD) (Souza et al., 2020) approaches, can greatly benefit structure-based rational drug design.

Rapid development of high-throughput screening using CRISPR/Cas9 system has greatly accelerated the discovery of new cancer drug targets in recent years (Fellmann et al., 2017; Jost and Weissman, 2018). CRISPR screening with tiling-sgRNA designs can further infer essential protein domains that are suitable for drug targeting (He et al., 2019; Neggers et al., 2018; Shi et al., 2015). However, identifying effective small molecular drugs for a specific target through high-throughput experimental screens is still expensive and inefficient (Macarron et al., 2011). Virtual screening using computational approaches has emerged as a starting point for identifying hit molecules for a given drug target (Lavecchia and Di Giovanni, 2013). 3D-based predictions of small molecules for a specific protein target with machine or deep learning approaches are of higher accuracy compared to sequence-based predictors. However, these ligand-based methods rely on multiple 3D structures binding the ligands to learn the features, thus are limited to a small fraction of ligands. Our study showed that conserved 3D binding patterns can be obtained at FG level, which may expand the scope for ligand-binding prediction since many different ligands share same or similar FGs.

## Methods and Materials

### Construction of the Datasets

We collected all the protein-ligand complexes from BioLiP database, a semi-manually curated database for biologically relevant ligand-protein interactions (Yang et al., 2013). Proteins that only bind metal ions were excluded. For each ligand, we removed the redundant proteins with more than 50% sequence similarity using cd-hit (Li and Godzik, 2006). Only ligands with at least 5 protein structures were kept for further analysis, producing a dataset containing 11570 protein structures in complex with 233 ligands. The PDB codes for all the protein structures, as well as the information of all 233 ligands are available at https://github.com/MDhewei/AFTME/Datasets.

For binding affinity analysis, we retrieved experimentally determined binding affinities of protein-ligand complexes from latest version of PDBbind database (Liu et al., 2015), resulting in binding affinity data for 599 complexes covering 158 out of 233 ligands in the dataset. Notably, the corresponding affinity values of each retrieved complex we used were p*K*_*d*_, *pK*_*i*_ and *pIC*_50_. These values were subjected to a transformation from raw affinity *K*_*d*_, *K*_*i*_ and *IC*_50_ defined as follows:

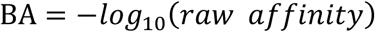

A higher value of BA indicates a stronger binding affinity for the protein-ligand complex.

### Ligand functional group definition and classification

Under the knowledge of biochemistry, we manually defined the functional groups (FGs) of each ligand based on their shape, size, and physicochemical property. Given a ligand in the dataset, we firstly downloaded its 2D structure from PDB database(Berman et al., 2000), then scanned its structure and search for FGs in the following order: (1) Ring structures with consistent physicochemical property like adenine; (2) Ring structures together with other polar groups like ribose; (3) Chain structures at the termini or in the middle of the ligand like alkane chain; (4) Well-defined polar groups such as phosphate, carboxyl, hydroxyl etc.; (5) Other fragments that are close in size with already defined FGs. In general, we followed a basic principal that the intraligand FGs should be considerably different in shape and physicochemical property but close in size, thus ensure their independency in interacting with the protein partner. For all the defined FGs, we further classified them into 21 major types referring to the classifications in previous study (Cai et al., 2008).

### AFTME Algorithm Description

AFTME takes four major steps to extract FG-binding motifs: 1. Extraction of ligand binding pockets. 2. Construction of functional atom distance matrix. 3.Two-dimensional clustering based on the distance matrix. 4. Identification and characterization of FG-binding motifs. Details of each step are described below.

#### Extraction of Ligand Binding Pockets

We defined the protein non-backbone heavy atoms within 5Å of the ligand as the functional atoms (FAs) (Hoffmann et al., 2010). And all FAs interacting with the ligand were gathered together to form the ligand binding pocket. To define the pocket described by a set of FAs that interact with the ligand, each FA consisting of the pocket with the bound ligand of length L was presented as a L-dimension vector:

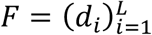

Where *d*_*i*_ represent the distance to the i-th atom from all L atoms of the ligand. Then the pocket P consisting of M FAs could be defined as follows:

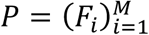

#### Construction of Functional Atom Distance Matrix

On completion of pocket definition, we constructed the functional atom distance matrix by collecting pockets binding the same ligand. Given K pockets that bind the same ligand of length L, we calculate the P vector for each pocket consisting M FAs with the vector F by extracting a set of FAs interacting with the ligand. For all of K pockets, we then obtained the matrix *M*_*FAD*_ as follows:

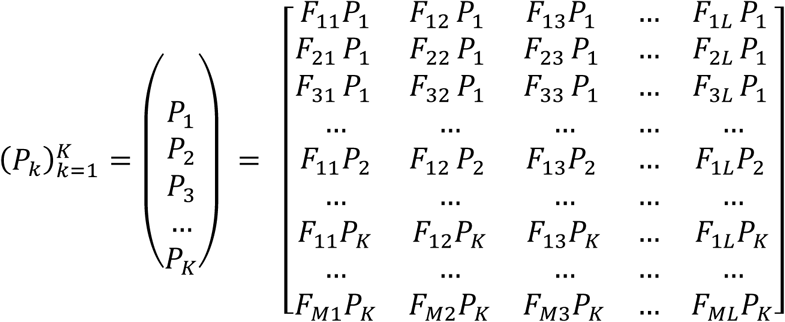

Where *F*_*ml*_*P*_*k*_ denotes the distance between *m-th* FA in the *k-th* pockets and the *L-th* atom of the bound ligand. Note M here just represents the number of FAs consisting of a pocket, it is not always equal for every pocket binding the same ligand. For example, considering 2 pockets binding same ligand with the length L, and we could extract the x, y FAs from that 2 pockets respectively. According to the definition of atom vector F and pocket vector P, we then calculated the *M*_*FAD*_ with the shape of x + y rows and L columns as follows:

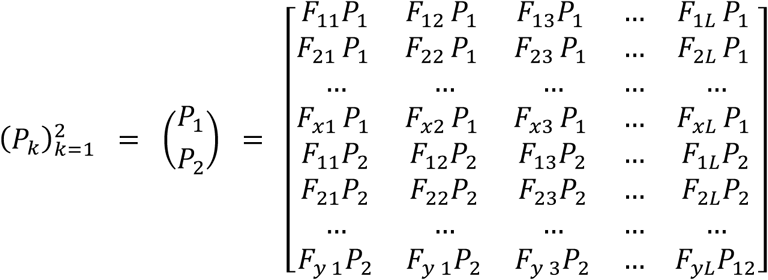

#### Two-dimensional clustering based on the distance matrix

Based on the functional atom distance matrix (*M*_*FAD*_) calculated for a specific ligand, we performed a two-dimensional hierarchical clustering both on the rows and columns of distance matrix, which represented the FAs and the ligand atoms (LAs) respectively. Prior to this analysis, we standardized the data within the rows of the matrix, i.e., subtract the minimum and divide each by its maximum. First of all, each FA/LA is considered as an individual cluster and the distances between different FA/LA were calculated through the Euclidean metric. Then two clusters with the closest distance were merged, and the linkages were created using the Ward method to minimize the total within-cluster variance. The clustering was performed using “AgglomerativeClustering” module from the scikit-learn package in python (Abraham et al., 2014). We set the “n_clusters” parameter as the number of predefined FGs of the ligand.

#### Visualization and identification of FG-binding motifs

We used a heatmap with a row-oriented and a column-oriented dendrograms to visualize the hierarchical clustering results. Based on the heatmap, we could find the correspondence between different cluster of FAs and LAs. Specifically, the observation that the cluster of FAs to its proximal cluster of LAs should clustered together, thus build up a map between different FA clusters and LA clusters. The clusters of LAs denote the different FGs within the ligand, and the clusters of FAs represent binding motifs for the specific FG matched. Following this step, we obtained different binding motifs for the FGs of the ligand they interact with. To maintain the consistency of LA clusters and our predefined FGs and reduce the noises in the identified binding motifs, we adopted a two-step filtering process: First, the LA clusters with less than 50% atoms in any of the predefined FGs were discarded together with the corresponding binding motifs, and vice versa. Second, protein pockets with less than 5 atoms were filtered within a specific binding motif.

#### Quantitative representation of FG-binding motifs

To quantitatively describe the identified binding motif, we made a deep insight into its composition from the protein level, amino acid level and atom level. In terms of protein and amino acid levels, we counted the number of binding pockets for all motifs interacting with specific FGs of the ligand, and the presence of each type of 20 amino acids inside a binding motif. In particular, atoms were classified into 6 categories according to their biochemical properties (He et al., 2015): (i) hydrophilic, (ii) acceptor, (iii) donor, (iv) hydrophobic, (v) aromatic, (vi) neutral. By calculating frequency of occurrence for each atom category within the motif, the motif could be expressed from the perspective of atom level. With deception of three levels ranging from low to high resolution, we could gain a well-rounded and detailed understanding of the binding motif. To make a quantitative evaluation of the binding motif, we expressed it using a 26-dimensional vector

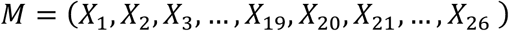

Where the former 20 dimensions are used to compute the proportion of occurrence of the amino acid *aa* in a specific motif for each of the 20 types of amino acids, and the latter 6 dimensions are used for the calculation of the proportion of occurrence of the atom properties *pp* of each category in the same motif for each of the 6 categories defined above. Particularly, the value in each dimension could be defined as follows:

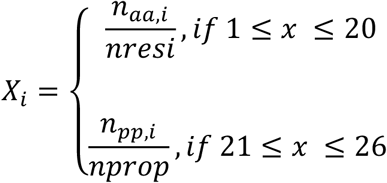

*n*_*aa,i*_ and *nresi* are the number of residues of type *i* amino acid *aa* observed in the motif and total number of residues in that motif, respectively. And the 20 types amino acids are assigned to a fixed order from 1 to 20. Similarly, *n*_*pp,i*_ and *nprop* are the number of properties of category *i* atom property *pp* and total number of properties, and the *i* ranging from 21 to 26 denotes the atom properties corresponding to the above 6 biochemical categories.

### Conservation evaluation of FG-binding motifs

To quantitatively assess the reusability of two FG-binding motifs, we calculated the Pearson correlation coefficient (PCC) between two 26-dimension vectors representing two binding motifs:

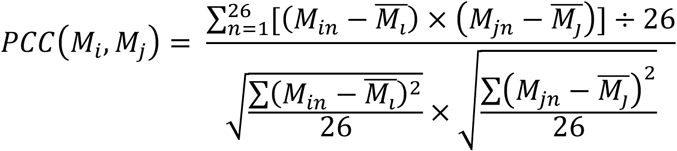

Higher PCC indicates stronger correlation between two binding motifs, which suggest their high reproducibility. To systematically measure the conservation of motifs binding same FG, we calculated the pair-wise PCC among all the motifs binding a specific FG, and evaluated the overall conservation score (CS) as the average of all the pair-wise PCC values:

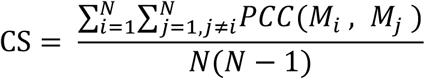

In addition, a permutation test was used to evaluate the statistical significance of the CS. Specifically, for each FG-binding motif appeared in multiple ligands, we randomly selected same number of motifs from all the identified motifs 1000000 times and calculated their corresponding CS value, the p-value was calculated as follows:

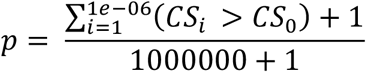

Where *CS*_*i*_ is the CS value of randomly selected motifs, *CS*_0_ is the CS value of motifs binding the same FG.

### Clustering on the 3D binding motifs

FG-binding motifs were classified based on their physicochemical properties, specifically, we performed k-means clustering on the 26-dimension vectors representing all the motifs. To determine the optimal number of clusters, we used elbow method which follows the basic idea to minimize the total intra-cluster variation as much as possible. Concretely, we first computed the k-means clustering on the data consisting of 481 vectors for different numbers of clusters k, which is ranging from 1 to 20. Next the total intra-cluster variation was calculated for each k value, and the formula is defined as follows:

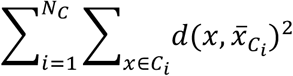

Where *N*_*C*_ is cluster numbers, *C*_*i*_ is the i-th cluster, *x̄* is the cluster centroid. Based on the computed variation under different values of k, a curve of the variation according to the number of clusters k could be plotted. Finally, the location of a bend (k=4) in the plot was selected as the optimal number of clusters in our approach.

## Availability of data and materials

The source codes of AFTME algorithm and the results of large-scale FG-motif analysis are available at https://github.com/MDhewei/AFTME. All other relevant data can be obtained from the authors upon request.

## Acknowledgements

This work was supported by National Natural Science Foundation of China (31621002 to L.N., U1632124 to L.N., and 31270770 to Z.Z.); Ministry of Science and Technology of China (2017YFA0504903 to L.N.); Hefei National Science Center Pilot Project Funds (in part). We thank all the lab members in Niu lab for helpful discussion.

## Author Contributions

W.H. and Z.L conceptualized the study. Y.L., W.H. and Y.Y. collected the data, Y.L. and

H.W. developed the method and performed the systematic analysis, Y.G., Z.Z. and M.K. helped data interpretation. L.N. and Z.L. supervised the project. Y.L., W.H., Z.L. and L.N. wrote the original manuscript. All authors read and approved the final manuscript.

