## Supplementary Figures for "Defining a Global Map of Functional Group Based 3D Ligand-binding Motifs"

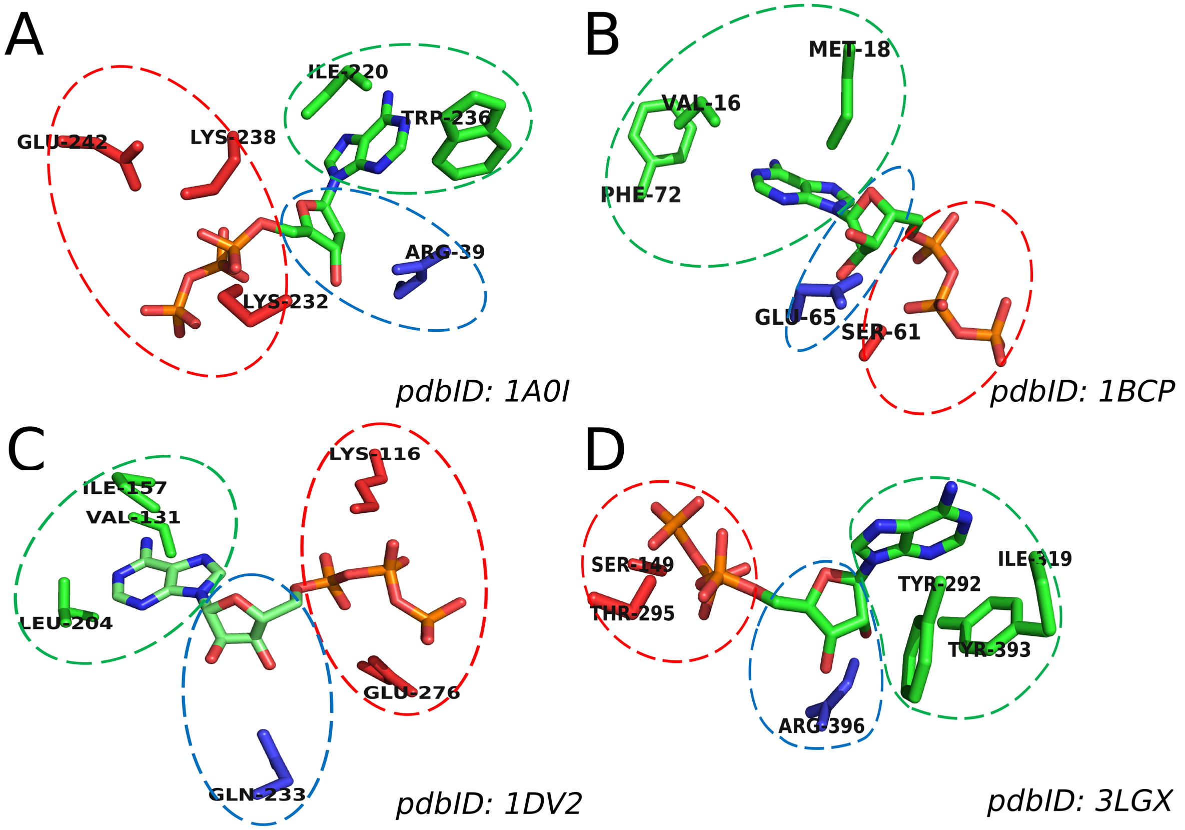


**Fig S1**: Four examples showing the 3D distribution of the amino acids within three different FG-binding motifs for ATP. Amino acids involved in different FG-binding motifs are marked in different colors: triphosphate-motif (red), adenine-motif (green), ribose-motif (blue), and are circled together with the corresponding FG using dash lines in the same color.

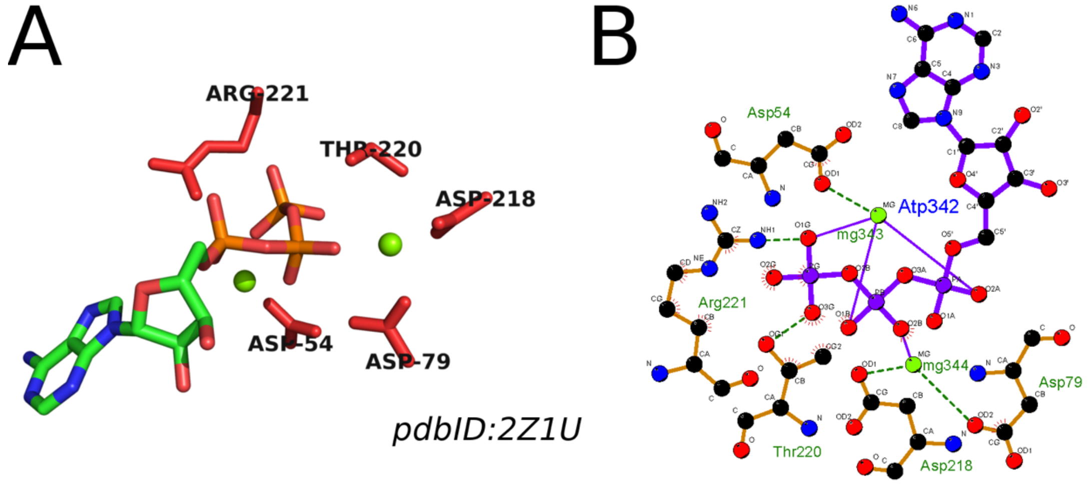


**Fig S2**: **The effect of metal ions in global ligand-binding profile.** (A). An example of ATP-binding sites composed of only motif for triphosphate group and two Mg ions. (B). The 2D protein-ligand interaction map showing how metal ions affect the global interaction patterns between functional atoms and ATP. The figure is generated with LigPlot software.

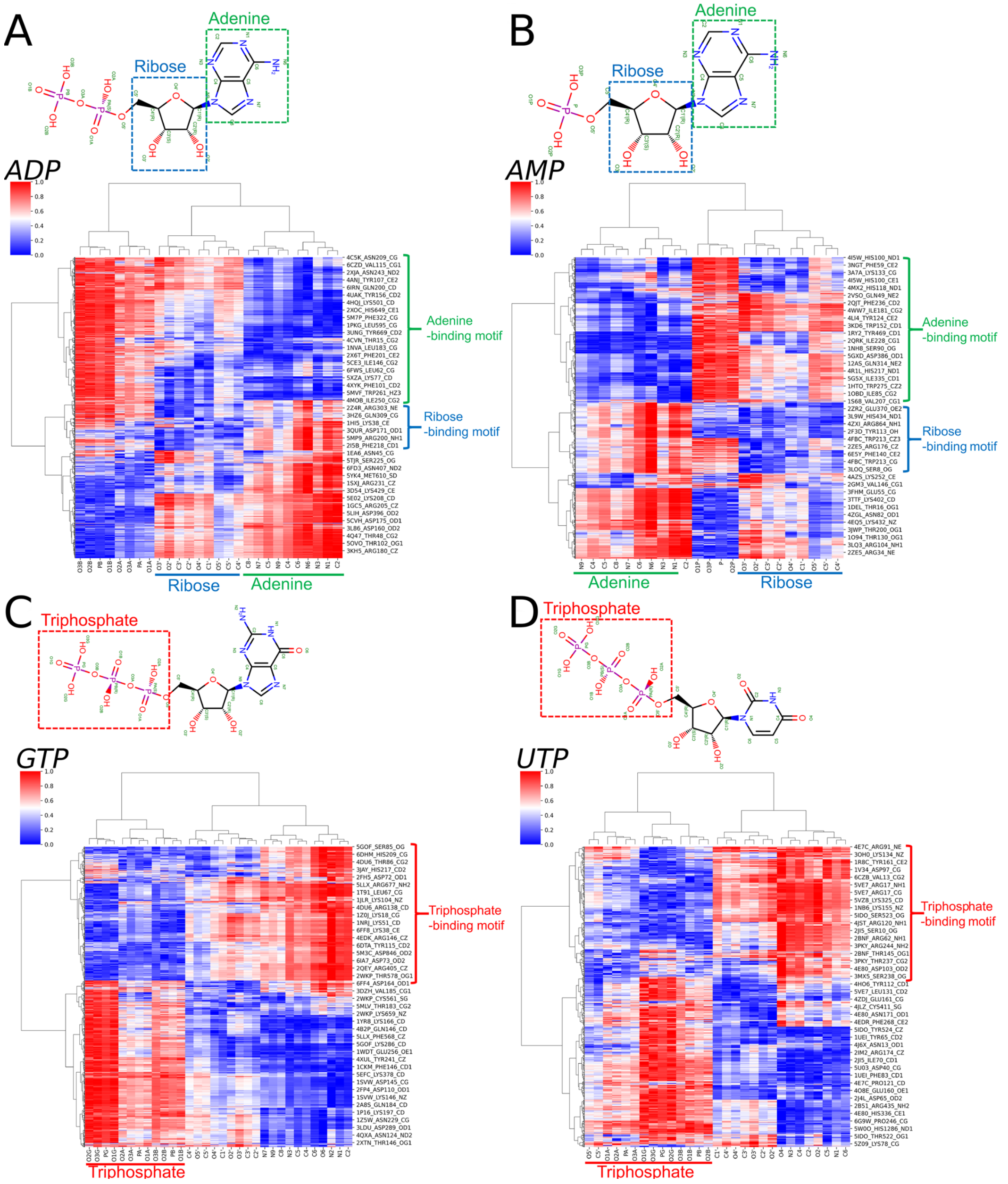


**Fig S3: FG-binding motifs for ligands share same FG with ATP**. (A-D). 2D structures and corresponding heatmaps showing the FG-binding motifs identified using AFTME for adenine and ribose in (A). ADP and (B). AMP, and triphosphate group in (C). GTP and (D). UTP. The functional group(s) shared with ATP in each ligand are marked with dashed rectangles.

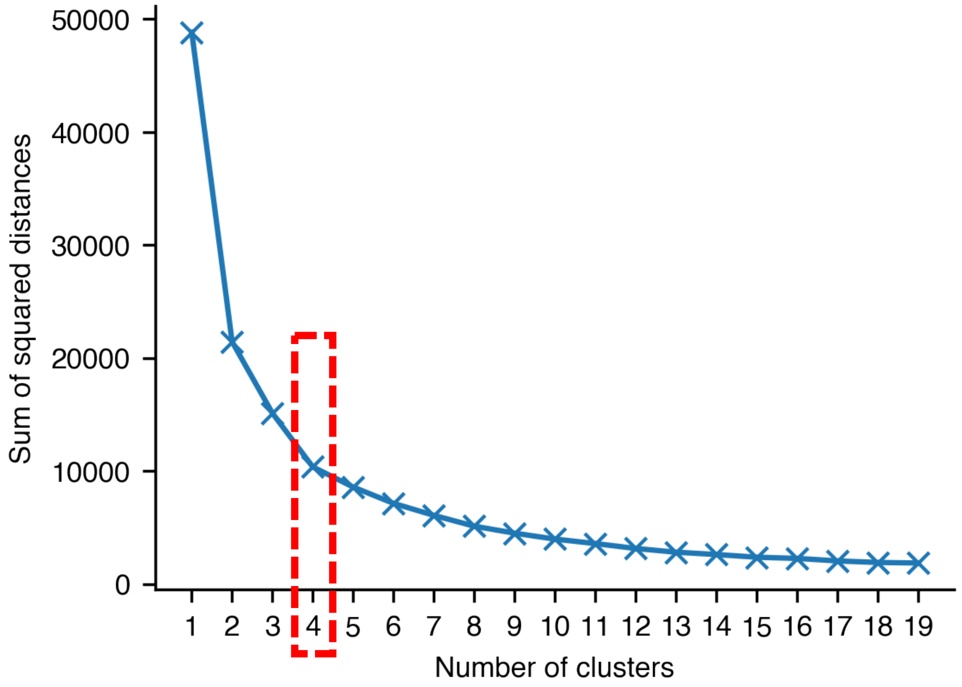


**Fig S4:** The plot showing the number of clusters selected in k-means clustering and the corresponding sum of squared distances. An optimal number of 4 was determined at the "elbow" of the plot.

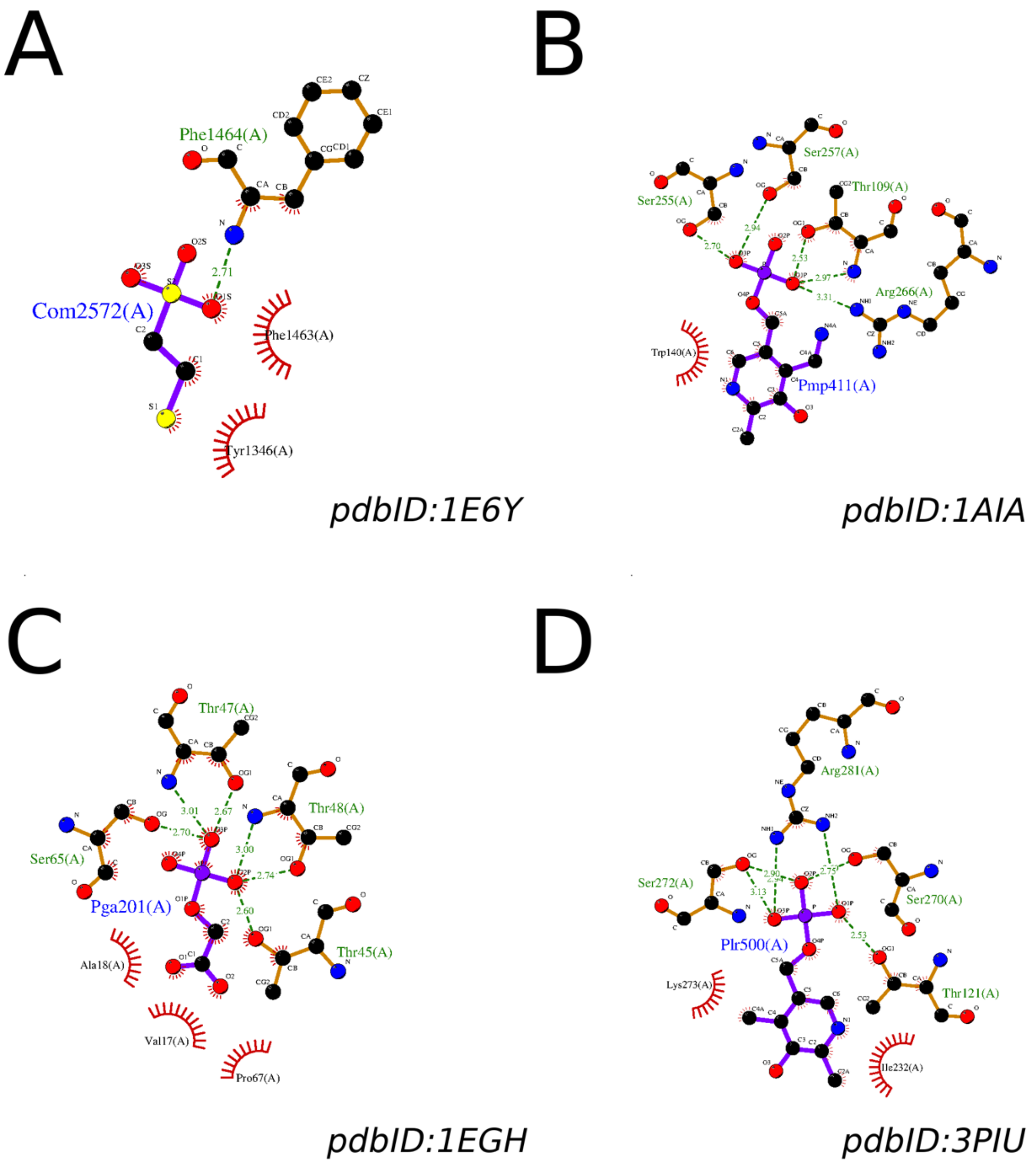


**Fig S5**: **Examples of hydrophobic-hydrophilic binding modes**. (A-D). 2D protein-ligand interaction maps for four different ligands contains a hydrophobic and a polar FG, which interact with hydrophobic and hydrophilic motifs respectively. The figures are generated with LigPlot software.

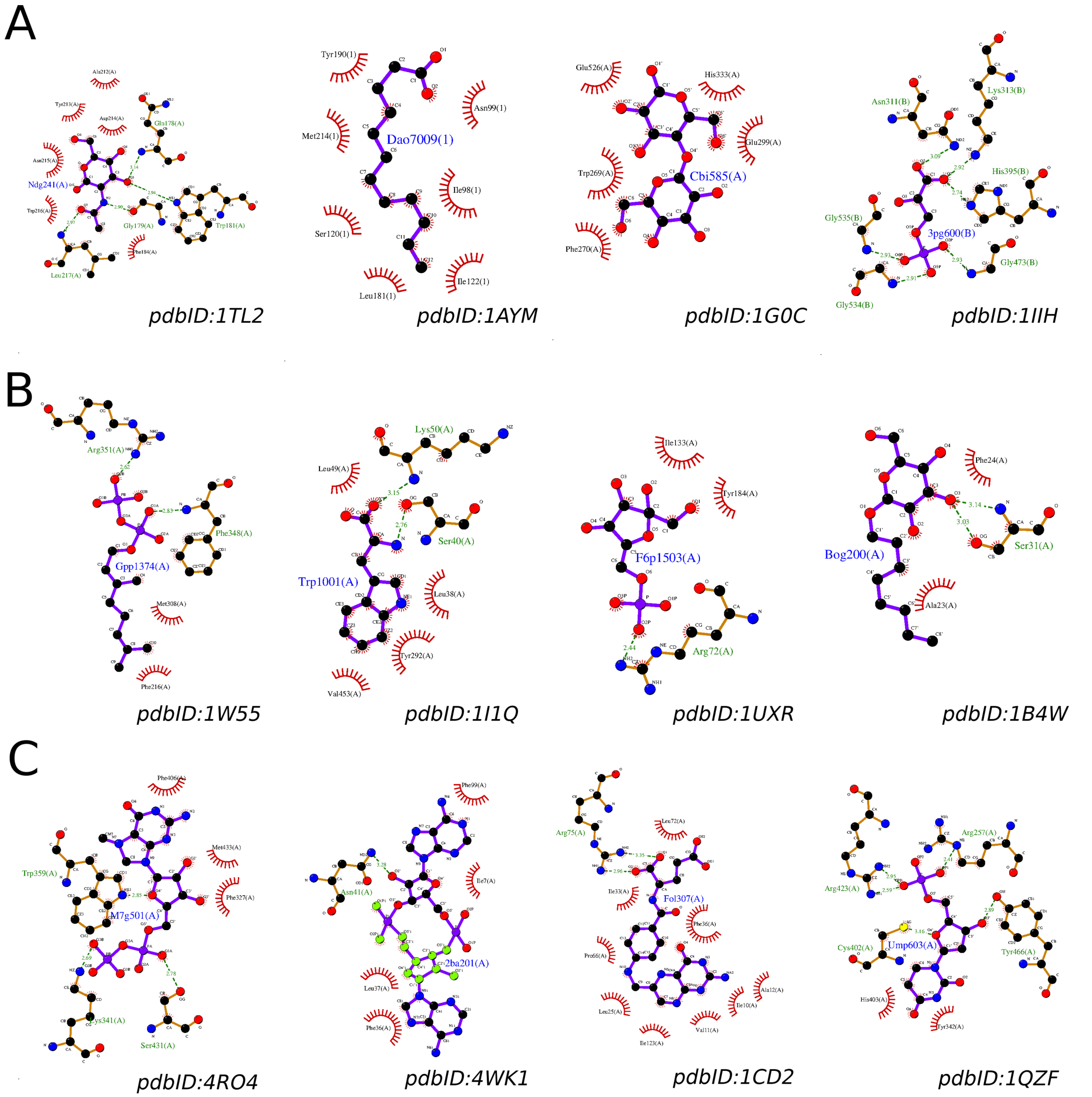


**Fig S6**: **Interaction patterns for different ligand binding modes**. (A-C). Examples of 2D protein-ligand interaction maps for different ligands involved in (A). same-type-binding mode, (B). two-type-binding mode and (C). three-type-binding mode. The figures are generated with LigPlot software.

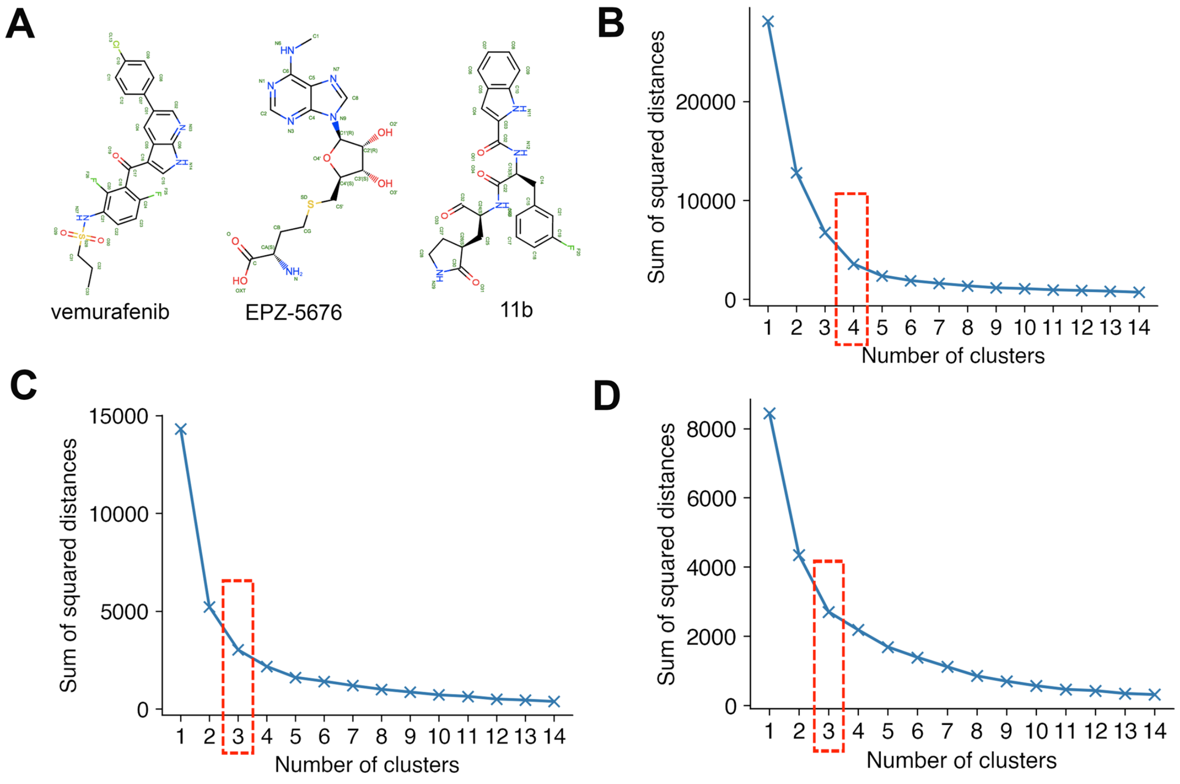


**Fig S7**: **Clustering of FAS for three small molecular drugs**. (A). The 2D structures of three small molecular drugs: the vemurafenib, the EPZ-5676 and 11b. (B-D). The plots showing the number of clusters against the corresponding sum of squared distances in k-means clustering for (B). BRAF-vemurafenib interactions (C). DOT1L-EPZ-5676 interactions and (D). Mpro-11b interactions. The optimal cluster number were selected at the "elbow" of the plots.
